# Structure of transcribing RNA polymerase II-nucleosome complex

**DOI:** 10.1101/437574

**Authors:** Lucas Farnung, Seychelle M. Vos, Patrick Cramer

## Abstract

Transcription of eukaryotic protein-coding genes requires passage of RNA polymerase II (Pol II) through chromatin. Pol II passage is impaired by nucleosomes and requires elongation factors that help Pol II to efficiently overcome the nucleosomal barrier^1-4^. How the Pol II machinery transcribes through a nucleosome remains unclear because structural studies have been limited to Pol II elongation complexes formed on DNA templates lacking nucleosomes^5^. Here we report the cryo-electron microscopy (cryo-EM) structure of transcribing Pol II from the yeast *Saccharomyces cerevisiae* engaged with a downstream nucleosome core particle (NCP) at an overall resolution of 4.4 Å with resolutions ranging from 4-6 Å in Pol II and 6-8 Å in the NCP. Pol II and the NCP adopt a defined orientation that could not be predicted from modelling. Pol II contacts DNA of the incoming NCP on both sides of the nucleosomal dyad with its domains ‘clamp head’ and ‘lobe’. Comparison of the Pol II-NCP structure to known structures of Pol II complexes reveals that the elongation factors TFIIS, DSIF, NELF, PAF1 complex, and SPT6 can be accommodated on the Pol II surface in the presence of the oriented nucleosome. Further structural comparisons show that the chromatin remodelling enzyme Chd1, which is also required for efficient Pol II passage^6,7^, could bind the oriented nucleosome with its motor domain. The DNA-binding region of Chd1 must however be released from DNA when Pol II approaches the nucleosome, and based on published data^8,9^ this is predicted to stimulate Chd1 activity and to facilitate Pol II passage. Our results provide a starting point for a mechanistic analysis of chromatin transcription.

---

To investigate how RNA polymerase II (Pol II) passes through chromatin, we determined the structure of Pol II transcribing into a nucleosome core particle (NCP). We initially attempted to obtain structures of elongating Pol II located at different positions within the NCP. To this end we placed transcription bubbles at different NCP locations, but these efforts were unsuccessful. We therefore reconstituted Pol II elongation complexes on pre-assembled, extended NCPs with different lengths of extranucleosomal DNA. The extended NCP that enabled structure determination comprised 145 base pairs (bp) of DNA with the Widom 601 sequence, 15 bp of extranucleosomal DNA, a 9-nucleotide (nt) 3’-DNA overhang, and a 10 nt RNA transcript annealed to the DNA overhang (Extended Data Fig. 1a). Pol II can bind the terminal DNA-RNA hybrid duplex of the extended NCP, resulting in a Pol II-NCP complex.

To test whether the reconstituted Pol II-NCP complex could elongate RNA, we incubated the complex with nucleoside triphosphates and transcription factor (TF) IIS, which is known to support Pol II elongation on nucleosomal templates^4^. We observed RNA elongation (Extended Data Fig. 1b), demonstrating that the reconstituted complex was functional. RNA elongation paused at several positions and was almost completely blocked after addition of ~35 nt (Extended Data Fig. 1b). The block to RNA elongation likely corresponds to the known major pause site around superhelical location (SHL) −5 of the NCP^3^. When we repeated the RNA elongation assay without histones, we observed extension to the end of the template (not shown), demonstrating that the presence of the NCP impairs RNA extension in this system.

To prepare a sample for cryo-electron microscopy (cryo-EM), we incubated reconstituted Pol II-NCP complex with nucleoside triphosphates and subjected the resulting complexes to size exclusion chromatography. TFIIS was initially included but lost during complex purification (Extended Data Fig. 1c, d). The purified sample was analysed by cryo-EM (Methods). Analysis of the obtained cryo-EM single particle data revealed a well-defined Pol II-NCP complex that could be refined to an overall resolution of 4.4 Å. Multi-body refinement resulted in a Pol II body at a resolution of 4.3 Å and a NCP body at 6.9 Å (Extended Data Fig. 2). The resulting reconstruction clearly showed nucleic acid backbones and grooves, well-defined protein secondary structure within Pol II, and the histone octamer fold of the NCP (Extended Data Fig. 3; Supplementary Video 1).

We next placed crystal structures of the Pol II elongation complex^10^ (PDB code 3HOV) and an NCP containing Widom 601 DNA^11^ (PDB code 3LZ0) into the cryo-EM density. The structures were locally adjusted to fit the density, connecting DNA was modelled, and the stereochemistry was refined (Methods). Pol II was observed in the active, post-translocated state (Extended Data Fig. 3f). However, we did not observe additional states of the complex with Pol II transcribing into the NCP, despite extensive sorting of the particle images. It is likely these states are unstable and did not survive preparation of the cryo-EM grids.

The structure reveals transcribing Pol II in front of a downstream nucleosome (Fig. 1). The orientation of the NCP with respect to Pol II is defined, and differs from the predicted orientation. To predict the Pol II-NCP orientation, we positioned structures of Pol II and the NCP on the designed nucleic acid scaffold assuming that the connecting DNA adopts the canonical B-form conformation. In this model, the nucleosomal disc is rotated by ~90° compared to the orientation observed in our experimental structure (Extended Data Fig. 4a). Comparison of the model with our structure also reveals that half a turn of nucleosomal DNA has been detached from the histone octamer surface. In particular, DNA base pairs between SHL −7 and SHL −6.5 are lifted from the octamer by up to 5 Å, and this enables feeding of the DNA into the polymerase active site cleft.

**Figure 1.**
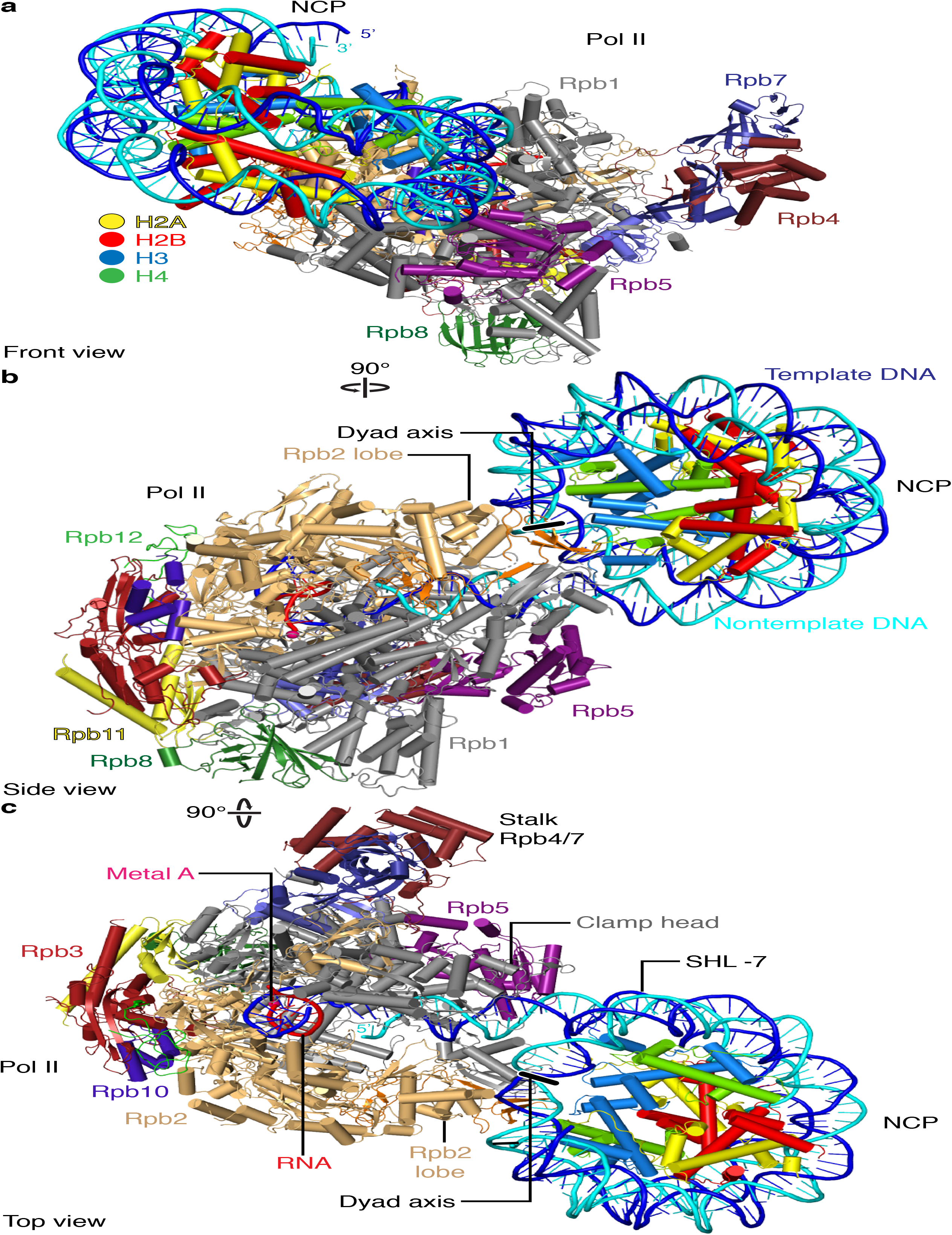
Structure of Pol II-NCP complex. **a-c**, Cartoon model viewed from the front (**a**), side (**b**), and top (**c**). Pol II subunits Rpb1, Rpb2, Rpb3, Rpb4, Rpb5, Rpb6, Rpb7, Rpb8, Rpb9, Rpb10, Rpb11, Rpb12, template DNA, non-template DNA, RNA, H2A, H2B, H3, and H4 are coloured in silver, sand, ruby, deep purple, slate, cyan, deepblue, forest green, orange, purple, yellow, green, blue, cyan, red, yellow, crimson, light blue, and green, respectively. Colour code used throughout. Histone octamer dyad axis is indicated as a black line with white outline.

These observations indicate that the defined orientation of the NCP results from Pol II-NCP contacts. Indeed, the structure reveals two major contacts between Pol II and the nucleosome (Fig. 2). Two Pol II domains that flank the polymerase cleft contact the NCP around its dyad axis. The clamp head domain in Pol II subunit Rpb1 contacts nucleosomal DNA around SHL +0.5, whereas the lobe domain of Rpb2 forms contacts around SHL −0.5. Pol II-NCP contacts involve the loops β4-β5 (Rpb1 residues 187-194) and α8-α9 (Rpb2 residues 333-345) that protrude from the clamp head and lobe, respectively.

**Figure 2.**
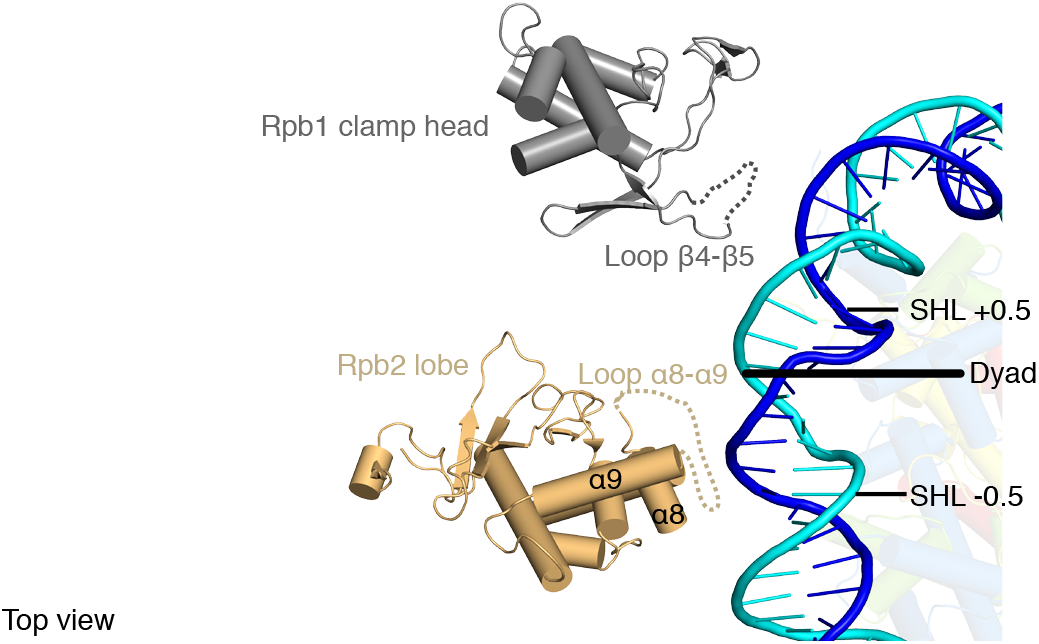
Pol II-NCP contacts. Rpb1 clamp head and Rpb2 lobe domains contact nucleosomal DNA around the NCP dyad. Clamp head and lobe domains are shown. Loops contacting nucleosomal DNA are indicated.

At the beginning of genes, Pol II often pauses. The sites of Pol II pausing are often found upstream of the first nucleosome in the promoter-proximal region (‘+1 nucleosome’). Pausing requires the elongation factors DSIF and NELF, raising the question whether these factors can be accommodated in the presence of the oriented nucleosome. Indeed, superposition of the Pol II-NCP structure onto our recent paused elongation complex structure (PEC)^12^ reveals that DSIF and NELF can be accommodated on the Pol II surface without clashes with the NCP (Fig. 3a). The resulting model of paused Pol II located in front of the +1 nucleosome reflects the situation *in vivo*, as follows. In the fruit fly *Drosophila melanogaster*, the +1 nucleosome adopts a defined location 135-145 bps downstream of the transcription start site^13-15^. This places the upstream edge of the +1 nucleosome at ~60-70 bp downstream of the TSS. Provided that ~15 bp of DNA are accommodated in the Pol II cleft, the active site of paused Pol II would be located 45-55 bp downstream of the TSS, in good agreement with known sites for promoter-proximal pausing^15-17^.

**Figure 3.**
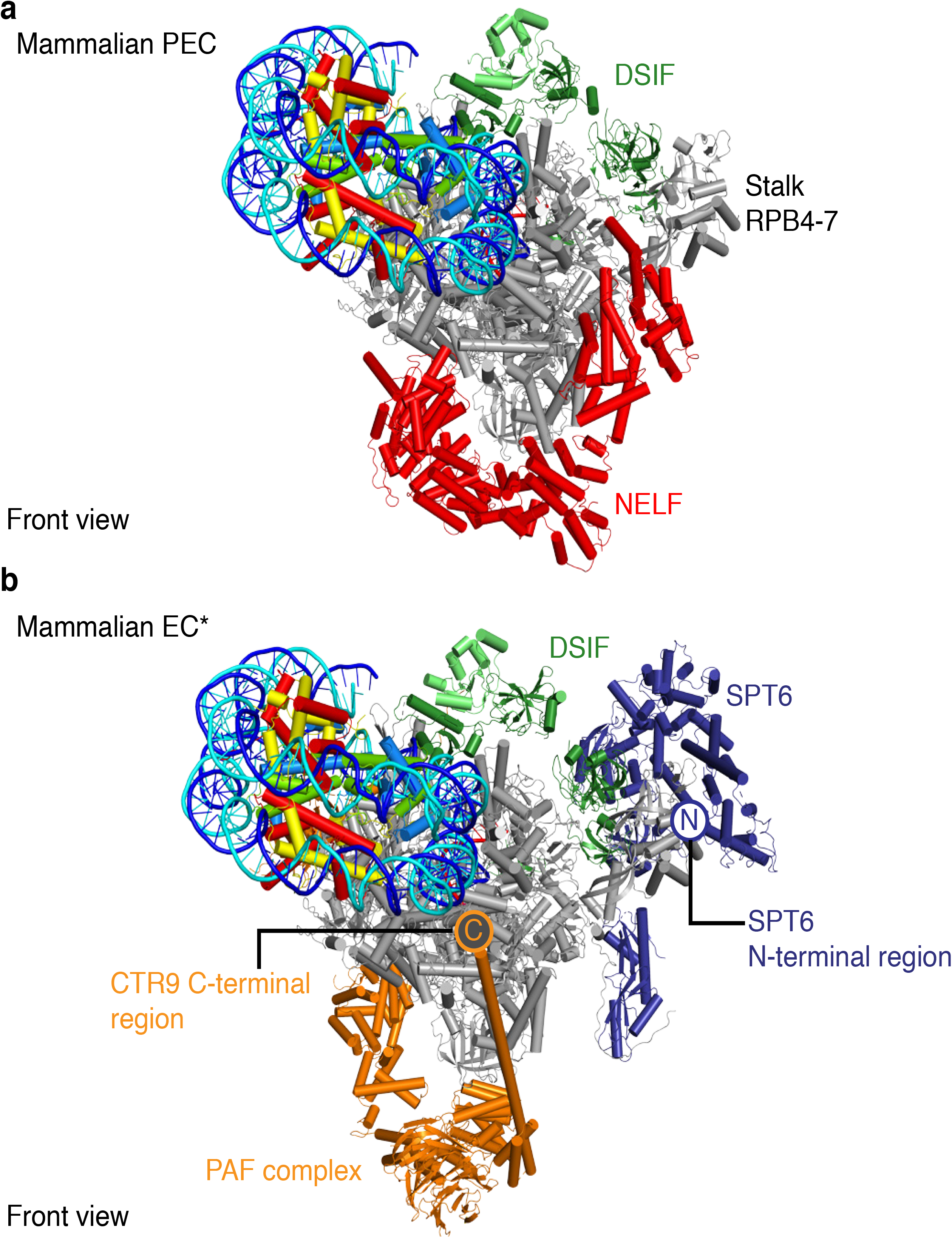
Accommodation of elongation factors. **a**, Superposition of Pol II-NCP structure with the mammalian paused elongation complex PEC (PDB code 6GML) provides the location of DSIF and NELF. **b**, Superposition of the mammalian activated elongation complex EC* (PDB code 6GMH) provides the location of PAF1 complex (PAF) and SPT6.

Release of Pol II from pausing and passage of Pol II through nucleosomes requires several protein factors, including TFIIS, the PAF1 complex (PAF), and SPT6^4, 18-22^. Superposition with our recent structure of the activated Pol II complex EC*^23^ shows that PAF and SPT6 can be accommodated in the presence of the NCP (Fig. 3b). In the resulting model, the C-terminus of the PAF subunit CTR9 is in close proximity to SHL +7 of nucleosomal DNA, explaining the known interaction of CTR9 with DNA^24^. The SPT6 N-terminal region is also located near the nucleosome, and is known to interact with the histone chaperone Spn1/IWS1^25,26^, which could bind histones when Pol II progresses and unravels nucleosomal DNA.

Nucleosome passage also requires the ATP-dependent chromatin remodelling enzyme Chd1^6,7^. Comparison of the Pol II-NCP structure with our recent structure of the Chd1-NCP complex^27^ after superposition of their NCPs revealed that the ATPase motor and the regulatory double chromodomain of Chd1 can be accommodated on the nucleosome surface without clashes with Pol II (Extended Data Fig. 4b). However, the DNA-binding region (DBR) of Chd1 would have to be displaced from its known position on extranucleosomal DNA^27,28^ when transcribing Pol II approaches the NCP. Previous studies showed that displacement of the DBR stimulates Chd1 ATPase activity^8,9^. Together these observations suggest that when Pol II approaches a Chd1-NCP complex, Pol II displaces the DBR of Chd1, thereby triggering full activation of the remodelling enzyme, which then loosens DNA-histone interactions. It however remains unclear how liberated histones will be transferred from downstream to upstream DNA, which involves the elongation factor FACT^1,29^.

It was reported that Pol II passage can lead to transient displacement of one of the two H2A-H2B dimers, converting the histone octamer to a hexamer^30^. The observed Pol II-NCP contacts suggest that the orientation of the NCP is maintained during further Pol II transcription. In this case, the proximal H2A-H2B dimer would encounter the clamp head and lobe domains when Pol II progresses by ~4-5 turns of DNA. The H2A-H2B dimer protrudes from the surface of the nucleosomal disc and would therefore clash with the lobe, possibly leading to its displacement. This model of nucleosome passage however raises the question how downstream DNA can be propelled into the active center cleft.

In conclusion, our structure of a transcribing Pol II-NCP complex provides a starting point for analysing the mechanisms of chromatin transcription. In the future, structures of Pol II-NCP complexes with elongation factors such as TFIIS, PAF, SPT6, IWS1, CHD1 and FACT should be determined. This should reveal how these factors assist Pol II in nucleosome passage. Ultimately, such studies could also show how the histone octamer is transferred from downstream to upstream DNA, to prevent loss of histones and to maintain histone modifications during transcription. We note that after our work was completed, others reported similar cryo-EM structures of Pol II-NCP complexes using a slightly different experimental strategy^31^.

## Acknowledgements

We thank past and present members of the Cramer laboratory. We thank C. Burzinski for purification of *S. cerevisiae* Pol II and TFIIS. S.M.V. was supported by an EMBO Long-Term Fellowship (ALTF-725-2014). P.C. was supported by the Deutsche Forschungsgemeinschaft (SFB1064, SFB860), the European Research Council Advanced Investigator Grant TRANSREGULON (grant agreement No. 693023), and the Volkswagen Foundation.

## Author Contributions

L.F. designed and carried out all experiments except biochemical experiments that were performed by L.F. and S.M.V.. P.C. designed and supervised research. L.F., S.M.V, and P.C. interpreted the data and wrote the manuscript.

## Author Information

Readers are welcome to comment on the online version of the paper.

## METHODS

No statistical methods were used to predetermine sample size. The experiments were not randomized, and the investigators were not blinded to allocation during experiments and outcome assessment.

### Protein preparation

*S. cerevisiae* Pol II was essentially purified as described^32^. *Xenopus laevis* histones were cloned, expressed, and purified as described^27,33,34^. *S. cerevisiae* TFIIS with an N-terminal GST tag was cloned, expressed, and purified as described^35^.

### Preparation of extended NCP

DNA fragments for nucleosome reconstitution were generated by PCR essentially as described^27,36^. A vector containing the Widom 601 sequence was used as a template for PCR. Large-scale PCR reactions were performed with two PCR primers (CGC TGT TTT CGA ATT TAC CCT TTA TGC GCC GGT ATT GAA CCA CGC TTA TGC CCA GCA TCG TTA ATC GAT GTA TAT ATC TGA CAC GTG CCT, reverse: ATC AGA ATC CCG GTG CCG AG) at a typical scale of 25 mL. The reaction was distributed into 48-well PCR plates (100 μl per well) and PCR was conducted with the following steps: 1. 98 °C for 1 min, 2. 98 °C for 10 s, 3. 72 °C for 45 s, cycle between steps 2 and 3 35 times, 4. 72 °C for 10 min, 5. Pause at 5 °C. PCR products were pooled, phenol chlorofom extracted, ethanol precipitated, and resuspended in 1 ml H_2_O buffer. The sample was ethanol precipitated, resuspended in 500 μl H_2_O buffer, and digested using TspRI (NEB) to generate the DNA 3’-overhang for RNA binding. Again, the digested DNA was phenol chlorofom extracted, ethanol precipitated and resuspended in 1 ml H_2_O buffer. The DNA was then applied to a ResourceQ 6 ml (GE Healthcare) and eluted with a gradient from 0–100% TE high salt buffer (10 mM Tris pH 8.0, 1 M NaCl, 1 mM EDTA pH 8.0) over a total volume of 60 mL at a flow rate of 2 mL/min. Peak fractions were analysed on a 1% (v/v) TAE agarose gel and fractions containing the DNA product were pooled, ethanol precipitated, resuspended in 100 μL H_2_O and stored at –20 °C. Preparation of extended NCP was performed as described^27^. Quantification of the reconstituted nucleosome was achieved by measuring absorbance at 280 nm. Molar extinction coefficients were determined for protein and nucleic acid components and were summed to yield a molar extinction coefficient for the reconstituted extended NCP.

### Reconstitution of Pol II-NCP complex

A RNA with the sequence /56-FAM/UCU CAC UGG A was purchased from Integrated DNA Technologies, resuspended in water, and stored at −80°C. To reconstitute the Pol II-NCP complex, Pol II, extended NCP and RNA were mixed at a molar ratio of 1:0.5:0.66 and incubated for 10 min at 30°C. Compensation buffer was added after 10 min to a final buffer of 150 mM KCl, 3 mM MgCl_2_, 20 mM Na⋅HEPES pH 7.5, 4% (v/v) glycerol, 1 mM DTT. For RNA extension, we added TFIIS (1:0.57, relative to Pol II) and NTPs (1 mM final concentration) and incubated for 30 min at 30 °C. The reaction was centrifuged (21,000*g*, 4 °C, 10 min), and applied to a Superose 6 Increase 3.2/300 column equilibrated in gel filtration buffer (100 mM NaCl, 3 mM MgCl_2_, 20 mM Na⋅HEPES pH 7.5, 5% (v/v) glycerol, 1 mM DTT). Peak fractions were analysed by SDS-PAGE and cross-linked with 0.1% (v/v) glutaraldehyde and incubated for 10 min on ice. The cross-linking reaction was quenched for 10 min using a concentration of 2 mM lysine and 8 mM aspartate. The sample was transferred to a Slide-A-Lyzer MINI Dialysis Unit 20,000 MWCO (Thermo Scientific), and dialysed for 6 h against 600 ml dialysis buffer (100 mM NaCl, 2 mM MgCl_2_, 20 mM Na⋅HEPES pH 7.4, 20 mM Tris⋅HCl pH 7.5, 1 mM DTT).

### RNA elongation assays

All concentrations refer to the final concentrations used in 70 μL reactions. Pol II (150 nM) was mixed with linear DNA or extended NCP (75 nM) and RNA primer (100 nM) at 30°C. Compensation buffer was then added to achieve final assay conditions of 150 mM KCl, 3 mM MgCl_2_, 20 mM Na⋅HEPES pH 7.5, 4% (v/v) glycerol, 1 mM DTT. TFIIS (90 nM) was added immediately before addition of NTPs. NTPs were added at a final concentration of 1 mM. 10 μL aliquots were taken at 0, 1, 5, 10, 20, and 30 minutes after NTP addition and were quenched by mixing the sample with 10 μL of Stop Buffer (6.4 M urea, 1x TBE, 50 mM EDTA pH 8.0). Reactions were treated with 4 μg proteinase K for 30 minutes at 30°C. The reactions were ethanol precipitated and resuspended in 5 μL of Stop Buffer and applied to a 7 M urea, 13.5% 19:1 Bis-acrylamide, 1x TBE sequencing gel run at 50W for 35 minutes in 1x TBE. RNA Products were visualized using the 6- FAM label and a Typhoon 9500 FLA Imager (GE Healthcare Life Sciences).

### Cryo-EM and image processing

The reconstituted and purified Pol II-NCP complex sample was applied to R2/2 gold grids (Quantifoil). The grids were glow-discharged for 100 s before sample application of 2 μl on each side of the grid. The sample was subsequently blotted for 8.5 s and vitrified by plunging into liquid ethane with a Vitrobot Mark IV (FEI Company) operated at 4 °C and 100% humidity. Cryo-EM data were acquired on a Titan Krios transmission electron microscope (FEI/Thermo) operated at 300 keV, equipped with a K2 summit direct detector (Gatan). Automated data acquisition was carried out using FEI EPU software at a nominal magnification of 130,000×. Image stacks of 40 frames were collected in counting mode over 9 s. The dose rate was 5.12 e^−^ per Å^2^ per s for a total dose of 46 e^−^ Å^−2^. A total of 9188 image stacks were collected.

Frames were stacked and subsequently processed. Micrographs were CTF and motion corrected using Warp^37^. Image processing was performed with RELION 3.0-beta 1^38^, unless noted otherwise. Post-processing of refined models was performed with automatic *B*-factor determination in RELION. Particles were picked using the neural network BoxNet2 of Warp, yielding 1,041,545 particle positions. Particles were extracted with a box size of 320^2^ pixel, and normalized. Particles were subdivided into three batches. Using a 30 Å low-pass filtered model from a previously published model (EMDB-3626), we performed iterative rounds of hierarchical 2D and 3D classification with image alignment and activated fast subset option (Extended Data Fig. 2c). 111,895 particles in batch 1 and 2 that showed Pol II-bound NCP were subjected to masked classification with the mask encompassing the NCP and global 2D classification. Batch 3 also showed Pol II-bound NCP and comprised 44,741 particles that were subjected to a masked 3D classification encompassing the Pol II-NCP complex. Particles from batch 1, 2, and 3 with NCP-bound Pol II were merged, 2D classified, and CTF refined^38^.

The final reconstruction was obtained from a 3D refinement with a mask that encompasses the Pol II-NCP complex. The Pol II-NCP reconstruction was obtained from 49,703 particles with an overall resolution of 4.4 Å (gold-standard Fourier shell correlation 0.143 criterion)^39^. The final map was sharpened with a *B*-factor of –182 Å^2^. Similarly to previous yeast Pol II cryo-EM structures, we observed an enrichment of certain Pol II orientations, exemplified in the angular distribution plot ^35,40^, but this did not compromise the resolution in any direction (Extended Data Fig. 2d, f). Local resolution estimates were determined using the built-in RELION tool. The Pol II-NCP complex was subsequently multi-body refined using two bodies encompassing Pol II or NCP with an estimated resolution of 4.3 Å for Pol II and 6.9 Å for NCP^41^ (Fourier shell correlation 0.143 criterion).

### Model building

Crystal structures of the *S. cerevisiae* Pol II EC (PDB code 3HOV) ^10^, and *X. laevis* nucleosome with Widom 601 sequence^11^ (PDB code 3LZ0), were placed into the density using UCSF Chimera^42^. Extranucleosomal DNA and nucleosomal DNA from SHL –7 to SHL –5 were built using COOT^43^. Secondary structure restraints were applied and the model was real-space refined against the post-processed EM map using PHENIX^44^. Figures were generated using PyMol^45^ and UCSF Chimera and ChimeraX^42^.

## EXTENDED DATA FIGURE LEGENDS

**Extended Data Figure 1.**
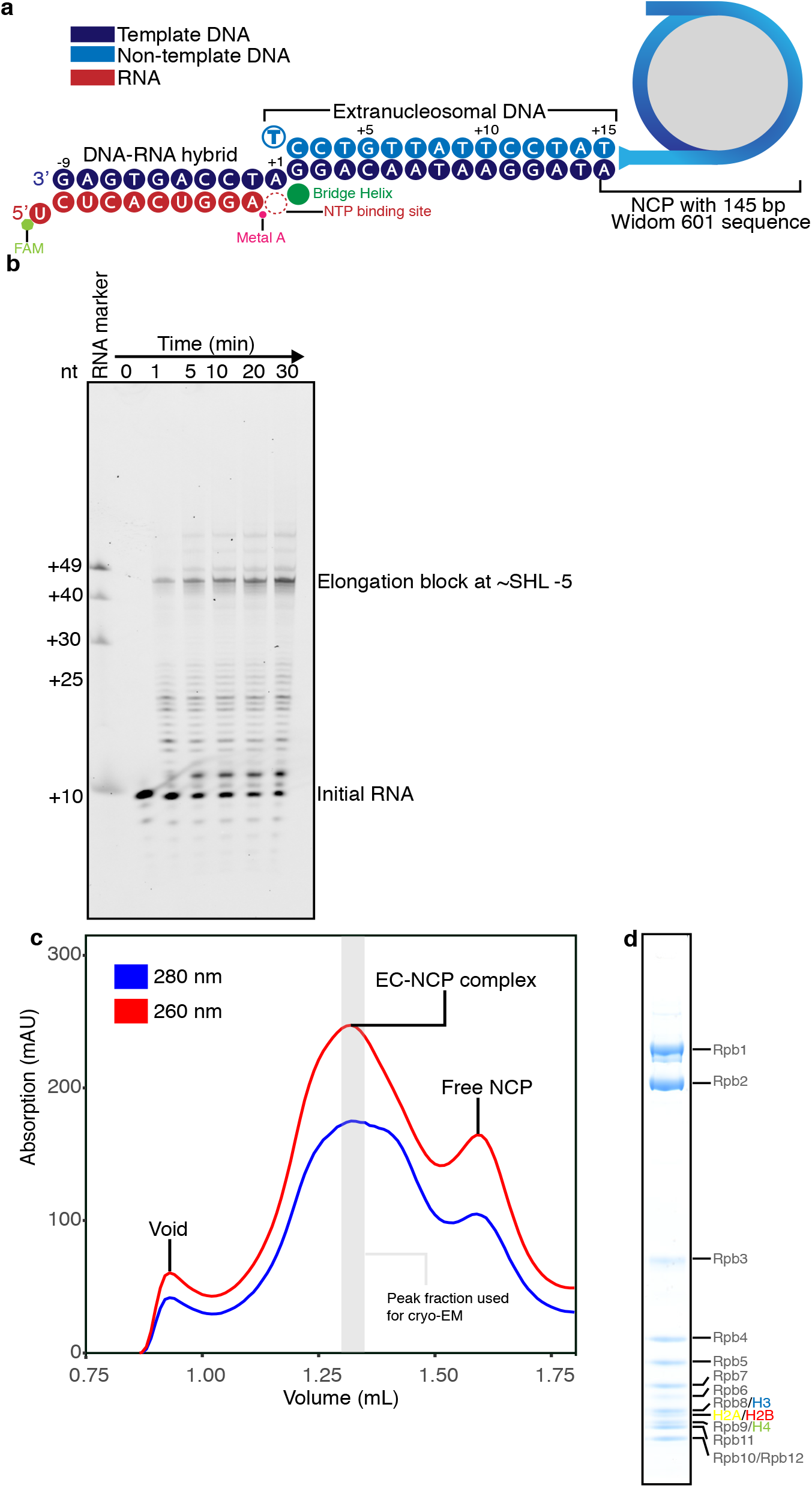
Formation of Pol II-NCP complex. **a**, Schematic of nucleic acid scaffold used for RNA elongation assays and complex reconstitution for cryo-EM analysis. **b**, RNA elongation assays performed with Pol II (150 nM), NCP or linear DNA (75 nM), TFIIS (90 nM) with a FAM labelled 10mer RNA primer. Reactions were quenched at various time points. **c**, Formation of the Pol II-NCP complex on a Superose 6 Increase 3.2/30 size exclusion chromatography column. Blue and red curve shows absorption at 280 nm and 260 nm milli absorption units, respectively. **d**, SDS–PAGE of peak fraction from gel filtration (c) used for cryo-EM grid preparation containing Pol II and histones. The identity of the bands was confirmed by mass spectrometry. All experiments were performed at least three times.

**Extended Data Figure 2.**
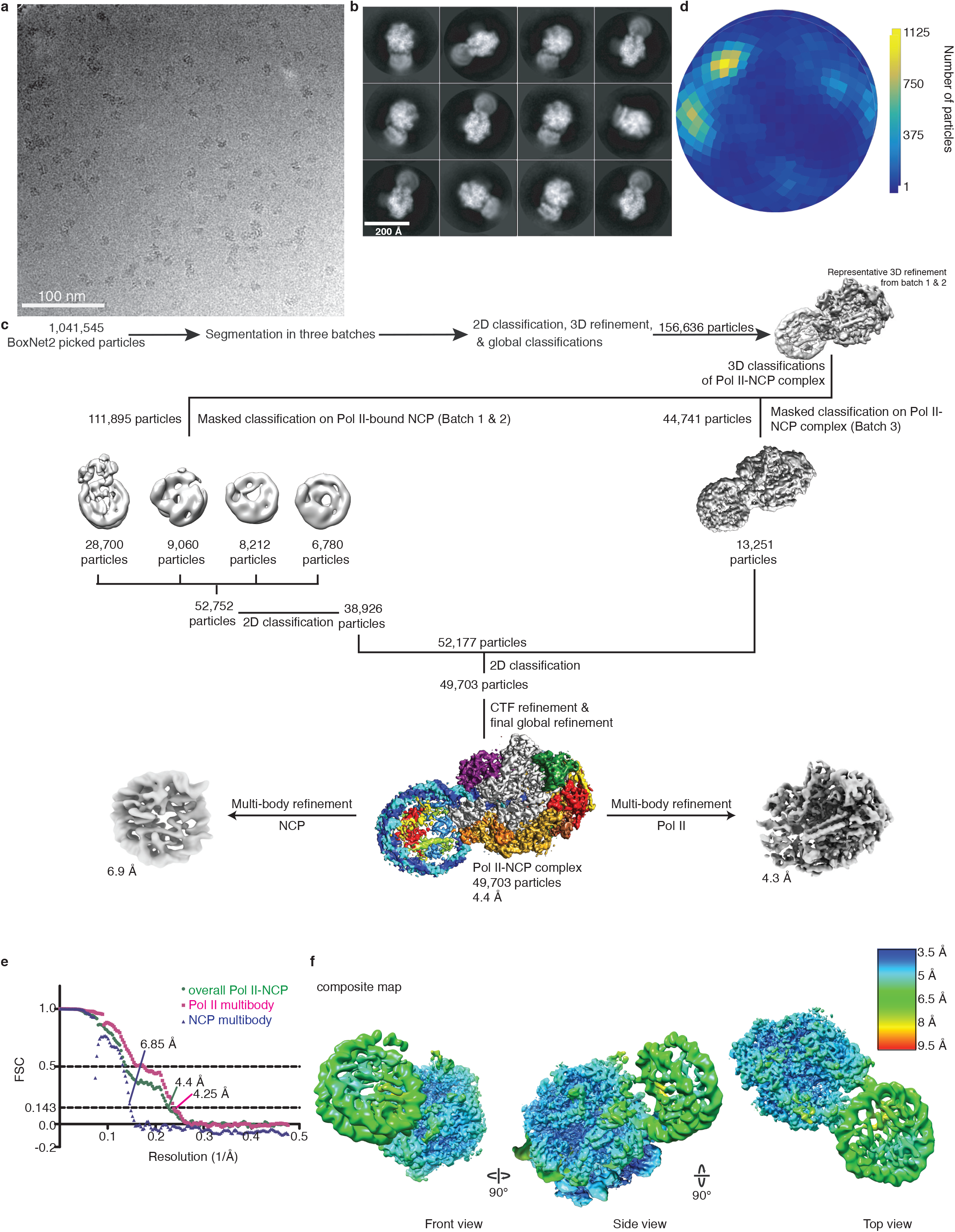
Cryo-EM structure determination. **a**, Representative micrograph, low pass filtered to enhance contrast. **b**, 2D class averages show Pol II in close proximity to a nucleosome. **c**, Angular distribution. Shading from blue to yellow indicates the number of particles at a given orientation. **c**, Sorting and classification tree used to reconstruct the Pol II-NCP complex at 4.4 Å resolution. **e**, Estimation of the resolution. The dark green line indicates the Fourier shell correlation between the half maps of the reconstruction. Resolution is given for the FSC 0.143. **f**, Local resolution estimation. Shading from red to blue indicates the local resolution according to the colour gradient. The multi-body refinement densities are shown. Absolute values are given.

**Extended Data Figure 3.**
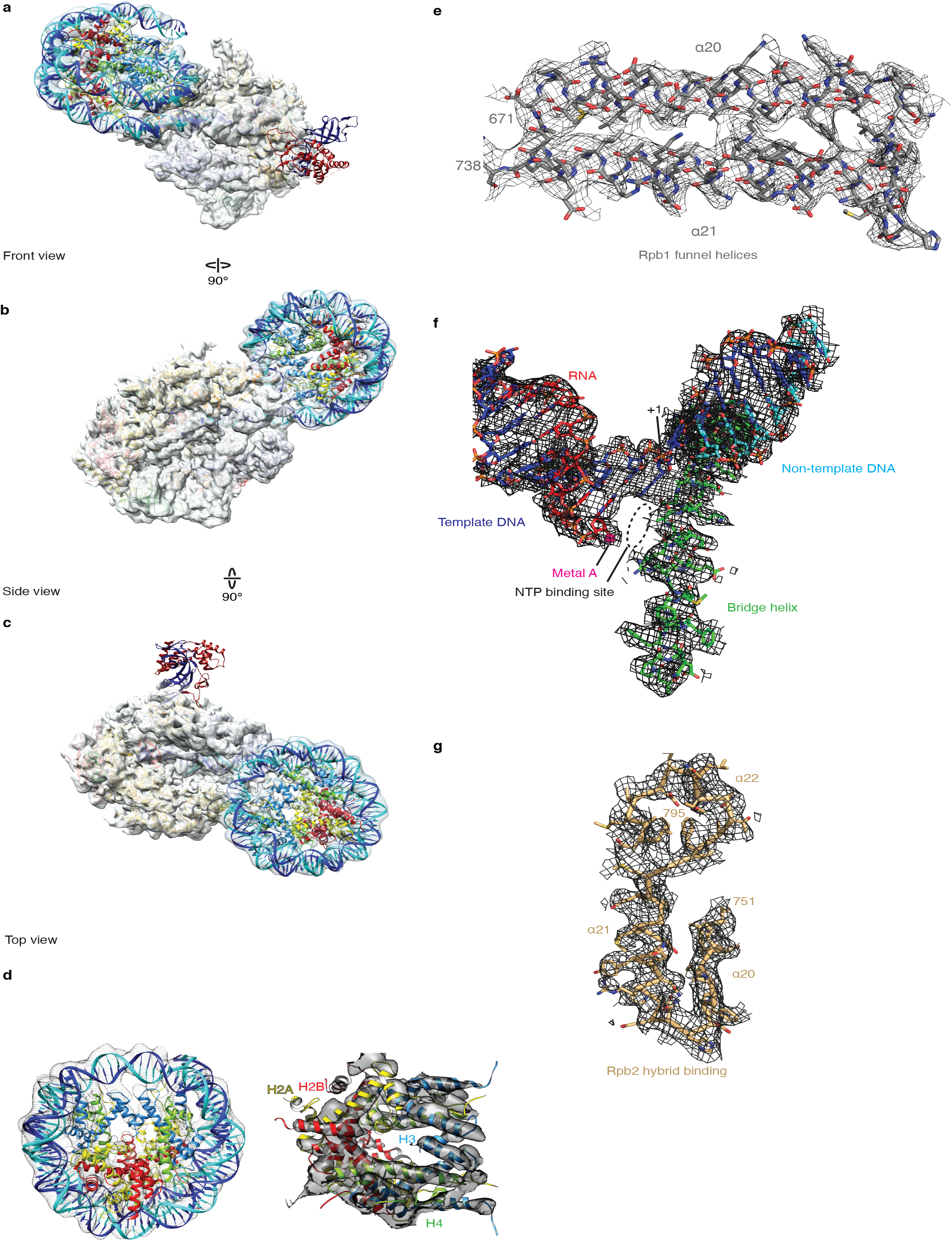
Cryo-EM densities. **a-c**, Non-sharpened maps after multibody refinement in (a) front, (b) side, and (c) top views. Pol II density is shown in silver, NCP density in light blue. Rpb4-7 is visible at lower contours (not shown). **d**, NCP density from multi-body refinement resolves the histone octamer fold and nucleosomal DNA. **e**, Rpb1 funnel helices on the Pol II surface are well defined. **f**, DNA-RNA hybrid in the Pol II active site adopts a post-translocated conformation with a free site for the nucleoside triphosphate substrate. **g**, Rpb2 hybrid binding region shows Pol II structure is very well resolved.

**Extended Data Figure 4.**
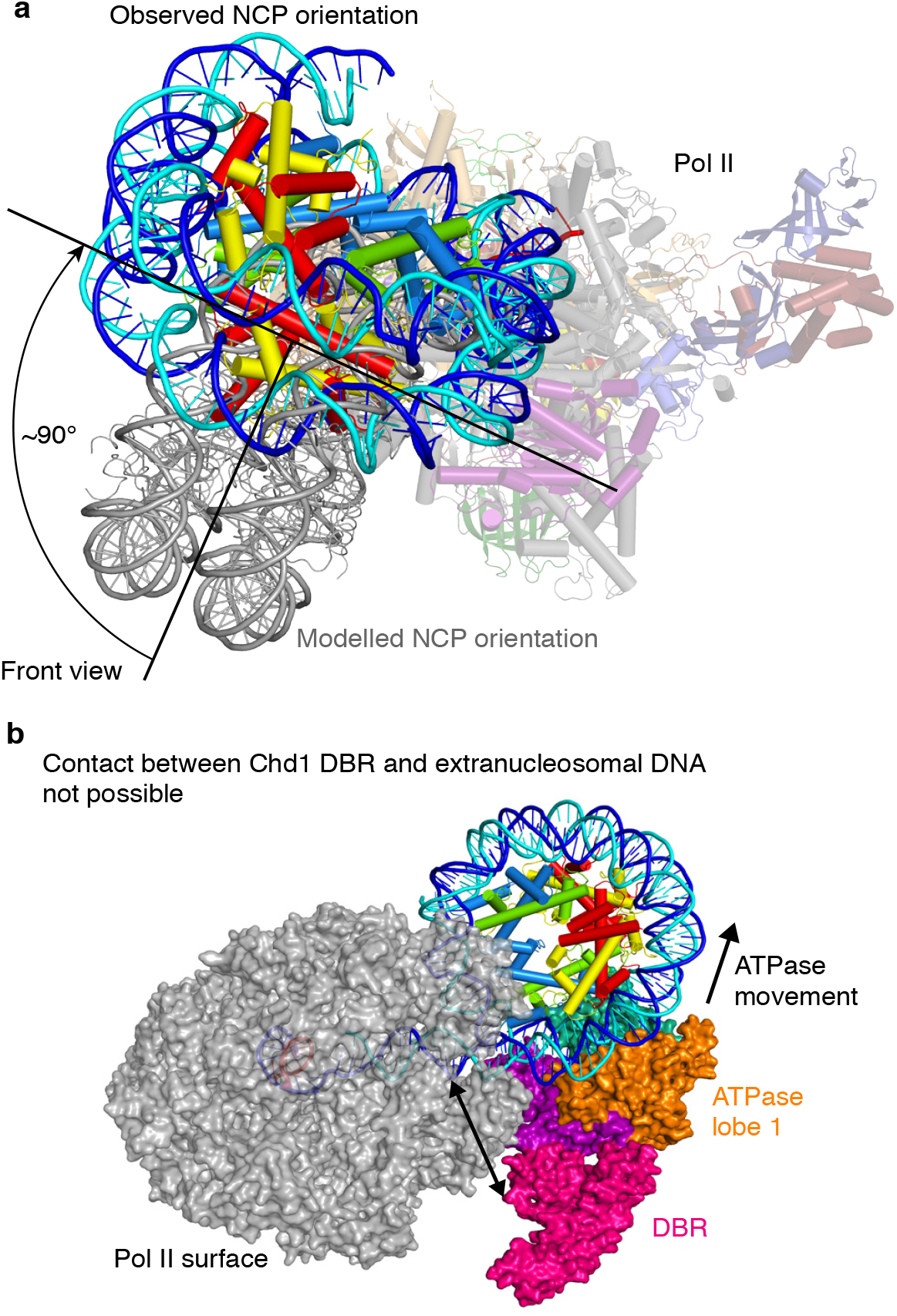
Additional models and structural comparisons. **a**, Downstream DNA of a Pol II elongation complex was modelled with a total length of 15 base pairs, followed by a mononucleosome (PDB code 3LZ0, shown in grey) based on our nucleic acid scaffold design. The experimental structure is coloured with the same colouring scheme used throughout the publication. Difference in rotational position of the NCP is indicated. **b**, Superposition with the Chd1-NCP complex (PDB code 5O9G) by aligning the NCPs reveals that the DNA-binding region (DBR) of Chd1 must be displaced from downstream DNA when Pol II approaches the NCP. Chd1 double chromodomain, ATPase lobe 1, ATPase lobe 2, and DNA-binding region are coloured in purple, orange, sea green, and pink, respectively.

**Extended Data Table 1.** Cryo-EM data collection, refinement and validation statistics.

|  | Pol II-NCP structure<br>(EMDB-XXXX)<br>(PDB XXXX) |
| --- | --- |
| **Data collection and processing** |  |
| Magnification | 130,000 |
| Voltage (kV) | 300 |
| Electron exposure (e-/Å<sup>2</sup>) | 46 |
| Defocus range (µm) | 0.25-4 |
| Pixel size (Å) | 1.05 |
| Symmetry imposed | C1 |
| Initial particle images (no.) | 1,041,545 |
| Final particle images (no.) | 49,703 |
| Map resolution (Å) | 4.4 |
| FSC threshold | 0.143 |
| Map resolution range (Å) | 4-12 |
| **Refinement** |  |
| Initial model used (PDB code) | 3HOV, 3LZ0 |
| Map sharpening *B* factor (Å<sup>2</sup>) | -181.9 |
| Model composition |  |
| Non-hydrogen atoms | 44,012 |
| Protein residues | 44,003 |
| Ligands | 9 |
| *B* factors (Å<sup>2</sup>) |  |
| Protein | 138.10 |
| Ligand | 100.64 |
| R.m.s. deviations |  |
| Bond lengths (Å) | 0.006 |
| Bond angles (°) | 1.20 |
| Validation |  |
| MolProbity score | 1.95 |
| Clashscore | 7.37 |
| Poor rotamers (%) | 0.37 |
| Ramachandran plot |  |
| Favored (%) | 90.01 |
| Allowed (%) | 9.9 |
| Disallowed (%) | 0.09 |

**Supplementary Video 1 | Cryo-EM density and structure of the Pol II-NCP complex**

